# Air temperature limits aerobic capacity and scope in active birds

**DOI:** 10.64898/2026.09.09.750338

**Authors:** Elana Rae Engert, Ammeli Nilsson, Fredrik Andreasson, Andreas Nord, Jan-Åke Nilsson

## Abstract

Thermal tolerance limits are an important determinant of species vulnerability to climate change. However, most of our knowledge of thermal tolerance pertains to resting animals, which could yield unrealistic predictions about how air temperature will affect animals when they are active. To gain a better understanding of thermoregulatory contstraints in active birds, we induced zebra finches in captivity to exercise continuously in air temperatures ranging from cool to warm (12-36 °C) while we simultaneously measured their metabolic rate, body temperature, and evaporative water loss. During activity, birds overheated in air temperatures well below their resting thermal tolerance limit, and aerobic scope was reduced by 22% at the warmest temperature. Although zebra finches showed a 8-fold increase in evaporative water loss during exercise in the warmest temperature, this was not sufficiently high to compensate for the substantial increase in heat produced during activity. This could explain why birds became hyperthermic and showed reductions in metabolic rate in the higher air temperatures. Thus, our results provide a mechanistic explanation for how heat could limit aerobic capacity and scope in active birds. These findings support growing evidence that increased air temperature can lead to sub-lethal fitness costs in free-living birds.

## Introduction

Knowledge of the thermal tolerance limits of animals is important for understanding the vulnerability of populations to climate change (Briscoe et al., 2023; Mitchell et al., 2018). While heat waves can cause mortality directly, prolonged heat exposure can also lead to missed opportunities and the accumulation of sublethal fitness costs which can ultimately lead to population declines (Conradie et al., 2019; Cunningham et al., 2021; Nord et al., 2026; Riddell et al., 2019). This has caused some to call for a more nuanced treatment of how thermoregulatory constraints are viewed in relation to ecology (Cunningham et al., 2021; Mitchell et al., 2018; Speakman & Król, 2010)

The heat balance of an animal in a given environment is maintained through adjustments to their heat inputs and outputs; metabolic heat production, dry heat loss, and evaporative heat loss (King & Farner, 1961; McNab, 1980). Endotherms maintain thermal equilibrium within a certain window of air temperatures while resting, referred to as the thermoneutral zone (TNZ), by modulating dry heat loss (Scholander et al., 1950). At temperatures below the TNZ, endotherms actively produce heat by increasing metabolic rate. When temperatures rise above the TNZ, they actively increase heat dissipation through evaporative cooling, for example by sweating or panting at the cost of water (McKechnie et al., 2021). In cooler temperatures, activity-generated heat can substitute for facultative thermogenesis. In warmer temperatures, hyperthermia will follow if metabolic heat production from activity is higher than the rate of heat dissipation.

When breeding in warmer temperatures, the capacity for heat dissipation can limit parental investment (Król & Speakman, 2003a, 2003b; Król et al., 2007). In birds, it has been shown that nestling provisioning rate is negatively correlated with environmental temperature on a global scale (Molenaar et al., 2026). While this represents compelling circumstantial evidence that increased environmental temperature constrains parental effort in birds, experimental evidence is needed to draw a direct link between air temperature, heat dissipation, and aerobic capacity. When heat dissipation has been experimentally facilitated by feather clipping, reproductive performance has typically improved via higher nestling provisioning rates (Tapper et al., 2020a), larger nestlings (Nord & Nilsson, 2019; Tapper et al., 2020b), reduced instances of hyperthermia (Nord & Nilsson, 2019; Tapper et al., 2020a), and lower parental mass-loss (Nord & Nilsson, 2019). Under some circumstances, such as when weather was cooler than usual, feather clipping has led to lower nestling provisioning rates than controls, presumably because the thermoregulatory costs for heat production increased, suggesting that heat substitution from activity did not completely offset heat loss (Tapper et al., 2020b).

The majority of work on thermal tolerance limits in birds and other endotherms has been performed on resting individuals, despite that reproduction depends on sustained activity. The scarcity of studies of active birds is likely explained by the difficulty of inducing sustained activity while measuring metabolic heat production (MHP), evaporative water loss (EWL), and body temperature (T_b_), while manipulating air temperature (T_a_). The studies that have accomplished some of these tasks have used the doubly-labeled water (DLW) method (Engel et al., 2006), mask respirometry (Nomoto et al., 1983; Tucker, 1968), mass-loss (Engel et al., 2006; Torre-Bueno, 1978), or thermal imaging (Lewden et al., 2023; Ward et al., 1999) in birds flying in wind tunnels or running on treadmills (Nomoto et al., 1983), but none have directly measured metabolic heat production, evaporative heat loss, and body temperature concurrently. This leaves unanswered questions as to what causes birds to reach their thermal tolerance limit during activity, at what temperature threshold this occurs, and how this compares to when they are resting.

The aim of this study was to improve our understanding of how thermoregulatory constraints act on active compared to resting birds. We therefore saught to quantify the how metabolic heat production and evaporative heat loss influence heat and mass balance during exercise and at rest in different tempertatures. To this end, we used a hop-flutter wheel (Engert et al., 2026) to elicit exercise in zebra finches (*Taeniopygia guttata*), which allowed us to continuously measure metabolic rate, body temperature, and evaporative water loss while manipulating air temperature ranging from cool to warm (10 - 40 °C). Because MHP increases with exercise, we expected that inflection points for evaporative cooling and hyperthermia would be shifted to cooler temperatures compared to when resting. As a result, we predicted that aerobic capacity would be limited by the capacity for heat dissipation above this threshold.

## Materials and methods

### Bird housing

Twenty-eight male zebra finches were transported from large outdoor aviaries to Lund University and were housed in an indoor facility with an ambient temperature of 22 °C and with a 12-hour light-dark cycle (07:00 to 19:00) for at least two weeks before measurements began. They were kept in six cages (120 x 84 x 100 cm) with four to six birds in each cage with unlimited access to food (tropical finch mix with vitamins and mineral supplements), drinking water, grit, cuttlebone and water for bathing. At least one week before measurements began, all birds were implanted intraperitoneally with a temperature sensitive PIT tag (L x W: 12 x 2.1 mm, LifeChip bio-thermo, Destron Fearing, DFW Airport, TX USA) weighing 0.26 g, 1 – 2% of the birds’ body mass, using the method described in Persson et al. (2024). All birds returned to normal behavior in their cages within 2 hours after the procedure.

### Thermoregulation in resting zebra finches

We measured metabolic rate of zebra finches overnight using flow-through respirometry to measure resting metabolic rate (RMR) at different ambient temperatures and basal metabolic rate (BMR) in thermoneutrality. Dry (Drierite, granular calcium sulfate anhydrous CaSO_4_, W. A. Hammond Drierite Company of Xenia) atmospheric air was pushed at a rate of 876.3 ± 144.4 mL min^-1^ (mean ± SD, standard temperature and pressure, dry [STPD]) into four respirometry chambers. The chambers were made of glass (1.2 liters, IKEA) with a plastic locking lid, covered with aluminium tape and spray painted matte black, with an inlet and an outlet for air. The flow rate for each chamber was measured using a 0-20 SLPM mass flow meter (Alicat Scientific, Tucson, AZ, USA). The excurrent air from the chambers was subsampled at 175.4 ± 21.5 ml min^.-1^ (mean ± SD, STPD), using an SS-4 Sub-Sampler Flow Meter and Pump (Sable Systems International). Water vapor pressure was measured using a RH300 Water Vapor Analyzer (Sable Systems) and then carbon dioxide CO_2_ using a CA-10 Carbon Dioxide Analyzer (Sable Systems). Then, water vapor and carbon dioxide were mechanically scrubbed from the air using Drierite and ascarite (II) (Acros Organics, Geel, Belgium) before oxygen O_2_ and barometric pressure (BP) was measured with an FC-10 Oxygen Analyzer (Sable systems). The CO2 sensor was zero and span calibrated using reference gases (100% N2 and 5% CO2, 95% N2) and the WVP sensor was zero calibrated using Drierite (Hammond Drierite Company, Xenia, OH, USA) immediately before the start of data collection. The O2 sensor was span calibrated with Drierite every morning before measurements began. Due to equipment malfunction, we could not use the data from the overnight recordings for two birds.

Four birds were taken at a time to the respirometry lab from the animal housing facility each evening. At 18:00, they were weighed before being separately placed into one of the four respirometry chambers, which were then placed inside a dark, climate-controlled cabinet (Weiss Umwelttechnik C180, Reiskirchen, Germany). The climate chamber was equipped with four antennas connected to HPR Plus data loggers Biomark, Boise, ID, USA) that recorded the body temperature of the four birds continuously. Air temperature inside each metabolic chamber was measured continuously using 36-gauge type T thermocouples and varied by ± 1 °C (SD) of the target temperature (Table S1). The birds acclimated in the climate cabinet for 1 h and 40 min at a temperature of 25 °C, similar to the air temperature in their housing, after which the temperature changed to 10 °C for the first RMR measurements. To measure RMR in different temperatures, we used a heat ramp protocol where the temperature was increased at 5 °C intervals between 10 - 40 °C (supplementary material, Figure S1), which includes temperatures both below and above the TNZ of zebra finches (32 – 39 °C, Briga and Verhulst, 2017). The temperature was held constant inside the climate chamber for a total of 40 min, during which time the four birds’ metabolic rate was measured consecutively for 10 min each. After this, the target temperature was increased by 5 °C which took about 20 minutes. During this time, baseline air measurements were taken. At 03:00, after the heat ramp measurements were completed, the temperature was decreased to 35 °C, a temperature within the TNZ, to record BMR for 5 hours. At 08:00, the birds were taken out of the chambers, weighed, and returned to their cages in the animal housing facility.

### Thermoregulation in active zebra finches

Each individual was measured in four temperature settings in the climate cabinet; 10, 20, 30 and 35 °C. One cage with up to six birds was measured per day at one of the set air temperatures. Between exercise measurements, there was a 10-13 day resting period, to allow birds to recover physically, and to prevent habituation. Each cage was tested in the four temperatures in a different order to balance all combinations of the order (1^st^, 2^nd^, 3^rd^, or 4^th^ session) and temperature (10, 20, 30, or 35 °C) to account for possible confounding effects of experience or habituation in the hop-flutter wheel. To ensure that each bird was tested at a similar time of day in all temperatures, the testing order of individuals in each cage was kept constant throughout the study.

To measure MHP, we used a hop-flutter wheel, which is a hermetically-sealed exercise wheel for birds (for details, see Engert et al., 2026). The respirometry setup was the same as the overnight recordings, but with an incurrent air flow rate of 4.46 ± 0.17 L min^-1^ STPD (mean ± SD) and the excurrent air was subsampled at a flow rate of 403.30 ± 57.00 mL min^-1^ STPD (mean ± SD). Baseline air was recorded for five minutes before and after each experiment. The temperature inside the wheel was continuously measured using a Thermichron iButton (OnSolution Pty Ltd., Castle Hill, Australia; 0.625 °C resolution; accuracy ± 0.5 °C) and was consistently 1 – 2 °C higher than the set temperature (Table S1). The four temperature settings will henceforth be referred to by the measured temperatures rounded to the nearest 1 °C. We used dry incurrent air and a high flow rate of 4.5 liters per minute to ensure that all metabolic measurements were conducted in low humidity conditions (< 1 kPa) to ensure that EWL was not constrained in exercising birds.

Starting at 9:00, birds were brought to the respirometry lab one at a time. At the start of each measurement, the bird was weighed and then placed inside of the wheel. We allowed birds to acclimate in the wheel for 30 minutes with the lights on and the glass window of the climate chamber covered to reduce disturbance, which was long enough for their body temperature to return to normal after experiencing stress-induced hyperthermia during handling. After the acclimation period, the window was uncovered to allow for continuous observation. The wheel started to spin at the lowest speed (0.1 m/s), which was slowly increased for 30 s until it reached 0.5 m/s, causing the bird to hop and flap. The speed of the wheel was adjusted continuously to maintain constant activity. The wheel was stopped when the bird could no longer control its position in the wheel, indicating that the bird was exhausted. The bird was allowed to rest and recover for 20 minutes in the wheel with the window covered, after which the bird was removed, weighed, and returned to its cage.

During some recordings, the bird was not induced to exercise and slid against the walls of the wheel instead of hopping and flapping. These recordings were excluded from the analysis. Some showed this behavior from the beginning and others developed it in the 2nd, 3rd or 4th trial, so some birds were not included in all four temperatures (Table S1). The birds’ behavior was not related to the temperature treatment (χ_2_ = 3.1, df = 3, p = 0.37).

### Data analyses

Respirometry data was processed and analyzed using ExpeData (v1.9.27) to extract the mean VO2 (mL min^-1^) during periods of interest. We used equation 1 (eqn 10.1 in Lighton, 2018) to calculate VO2, where flow rate (FR) is expressed in mL min^-1^, ΔO_2_ is difference in the fractional concentration of O_2_ from the baseline air and O_2_ is the fractional concentration of O_2_.

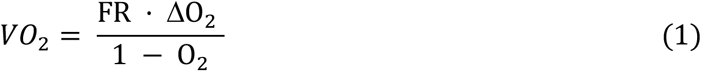

CO_2_ was corrected for water vapor dilution using equation 2 (eqn 8.7 in Lighton, 2018). VCO_2_ was calculated using equation 3, (eqn 10.6 in Lighton, 2018), where ΔCO_2_ is difference in the fractional concentration of CO_2_ from the baseline air, BP is barometric pressure, and WVP is water vapor pressure (kPa).

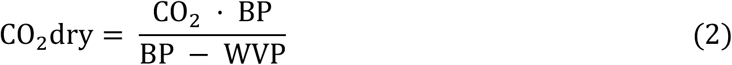

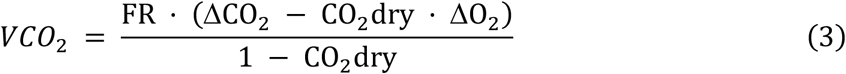

We converted to metabolic rate in Watts using equation 4 (based on eqn 9.13 in Lighton, 2018) where VO_2_ is the volume of oxygen in milliliters per minute and RQ is the individual respiratory quotient (VCO_2_ / VO_2_; mean ± SD: 0.86 ± 0.06).

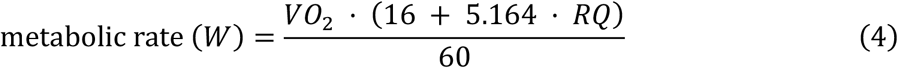

We calculated evaporative water loss (EWL, mg min^-1^) using equation 5 (based on eqn 10.9 in Lighton, 2018) by dividing WVP by the temperature of the air stream (T) temperature and the gas constant for water vapor (R_w_, 461.5 J kg^−1^ K^-1^) and multiplying the resulting water vapor density (WVD) by flow rate (FR, ml min^-1^).

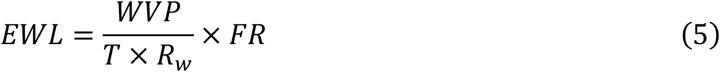

We calculated evaporative heat loss (EHL) using the heat of vaporization of water 2.406 J / mg H_2_O.

The mean RMR (resting metabolic rate) for each bird during overnight recordings was calculated as the most stable two minutes from the 10-minute recording during each five-degree interval from 10 - 40 °C. BMR was calculated as a mean from the most stable two minutes during the three 10-minute intervals when metabolic rate was measured (after the ambient temperature had been set to within the TNZ at 35 °C) and taken as the lowest out of those three means. From the exercise recordings, MHP was taken as a mean from the running five minutes period during exercise when oxygen consumption was highest. The mean body temperature during the overnight recordings was calculated from each two-minute period when RMR was measured. Body temperature during exercise was calculated as the mean of the five-minute period when MHP was measured.

For the overnight measurements of RMR at 10 – 40 °C, we used “broken-stick” mixed models with repeated measures, allowing for random individual slopes and intercepts, to identify break-points and slopes for the effect of T_a_ on resting metabolic rate (RMR), core body temperature (Tb), evaporative water loss (EWL), and evaporative cooling efficiency (EHL/MHP). We used RMR to estimate the lower critical temperature (LCT), so chamber temperatures above the expected upper critical temperature (UCT, 39 °C) were excluded prior to analysis. We restricted the search range for the break point to above 28 °C after visual examination of the plot of RMR and T_a_ to ensure that a realistic LCT was identified. The search range in all other models included the full range of data.

We investigated the effect of air temperature (T_a_ = 12, 22, 32, and 36°C) on MHP, metabolic scope, endurance, body temperature (T_b_), EWL, and evaporative cooling efficiency (EHL/MHP) using linear mixed effects models (LMMs). Body mass, the order of measurements, and time were included as fixed effects and ID was included as a random intercept in all models. Model residual plots were visually inspected to verify that the model assumptions were fulfilled. Endurance and EWL were log-transformed to fulfill the model assumptions. We used the *lme4* package to obtain model results (Bates et al., 2015). Estimated marginal means of categorical factors (temperature and order) and results of pairwise comparisons were obtained using the *emmeans* package with adjustments for multiple comparisons using the Tukey method (Lenth, 2022). We obtained the marginal and conditional R^2^ of LMMs (Nakagawa & Schielzeth, 2013) using the *performance* package (Lüdecke et al., 2021).

## Results

### Thermoregulation in resting zebra finches

For the lower critical temperature (LCT), we identified an inflection point in RMR at 33.9 °C (95% CI: 31.7 °C, 38.1 °C) (Table S2, Figure 1). The inflection point for Tb was estimated to be at 33.4 °C (95% CI: 32.8– 33.5 °C). The upper critical temperature (UCT), defined as the inflection point for EWL, was 36.8 °C (95% CI: 35.9 – 37.3 °C). Evaporative cooling efficiency (EHL/MHP) had a similar inflection point at 36.2 °C (95% CI: 33.8 – 36.9 °C).

**Figure 1.**
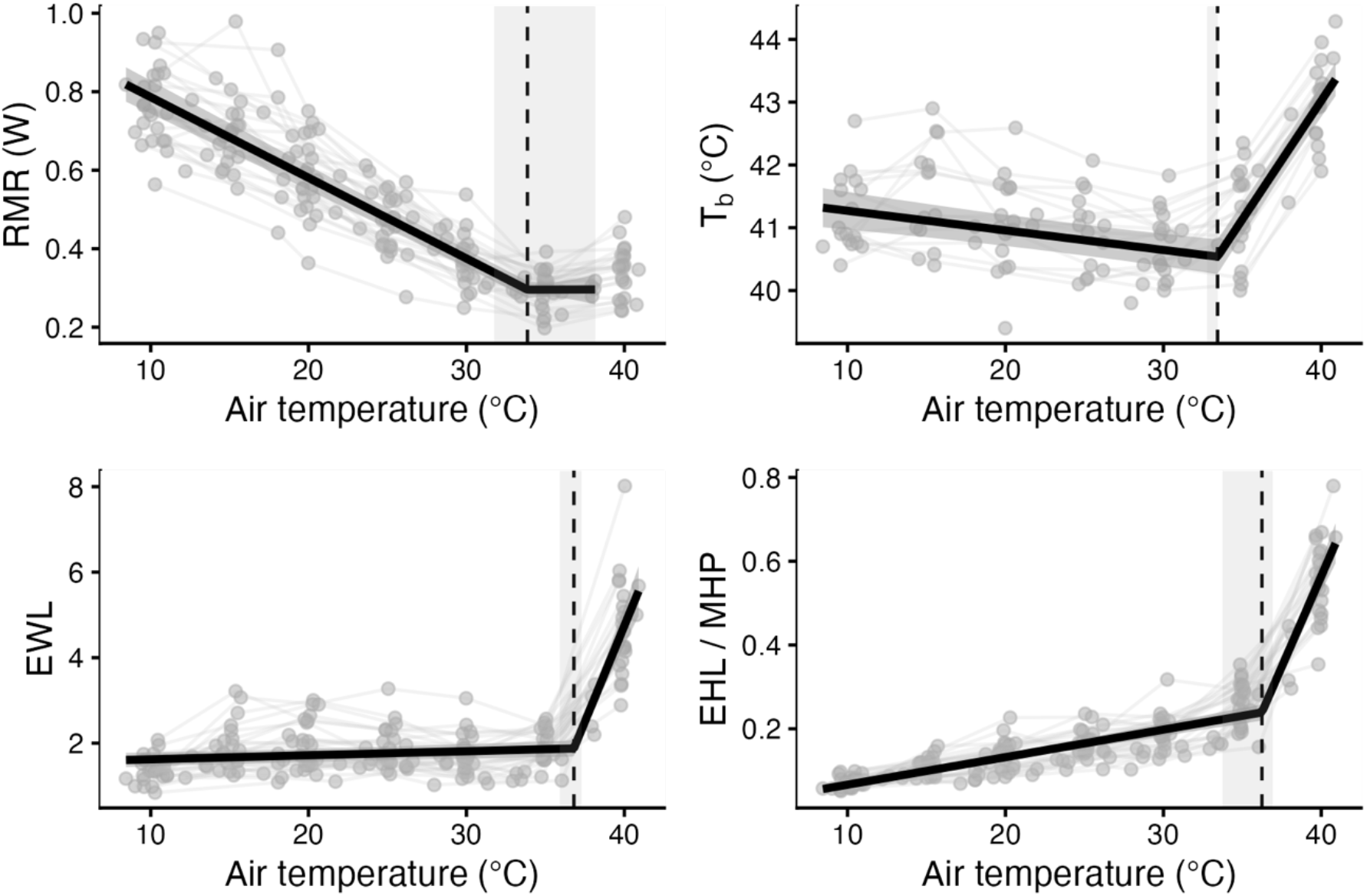
“Broken stick” mixed models with slopes ± 95% CI (solid black lines and gray ribbons) and breakpoints ± 95% CI (dashed lines and gray ribbons) for zebra finches measured overnight in different air temperatures.

### Thermoregulation in active zebra finches

MMR decreased with increasing air temperature. MMR was 18% lower at 36 °C compared to 12 °C and was 13% lower compared to 22 °C. The mean metabolic scope was 6.0 × BMR at 12 °C, and 4.7 × BMR at 36 °C, which was significantly lower than at 12, 22, and 32 °C. We did not find a significant effect of air temperature on endurance, but birds had the highest mean endurance of 22.3 minutes in 22 °C, and the lowest endurance of 14.6 minutes in 36 °C. T_b_ during MMR increased with increasing T_a_, reaching 45.1 °C when T_a_ = 36 °C. EWL and evaporative cooling efficiency (EHL/MHP) was significantly higher at 36 °C compared to all other temperatures, but did not differ significantly between 12, 22 and 36 °C (Figure 2, Table S3). Compared to pre-exercise, T_b_ decreased during MMR in 12 °C and increased in 32 and 36 °C (time point x air temperature interaction, Figure 3, Table S4).

**Figure 2.**
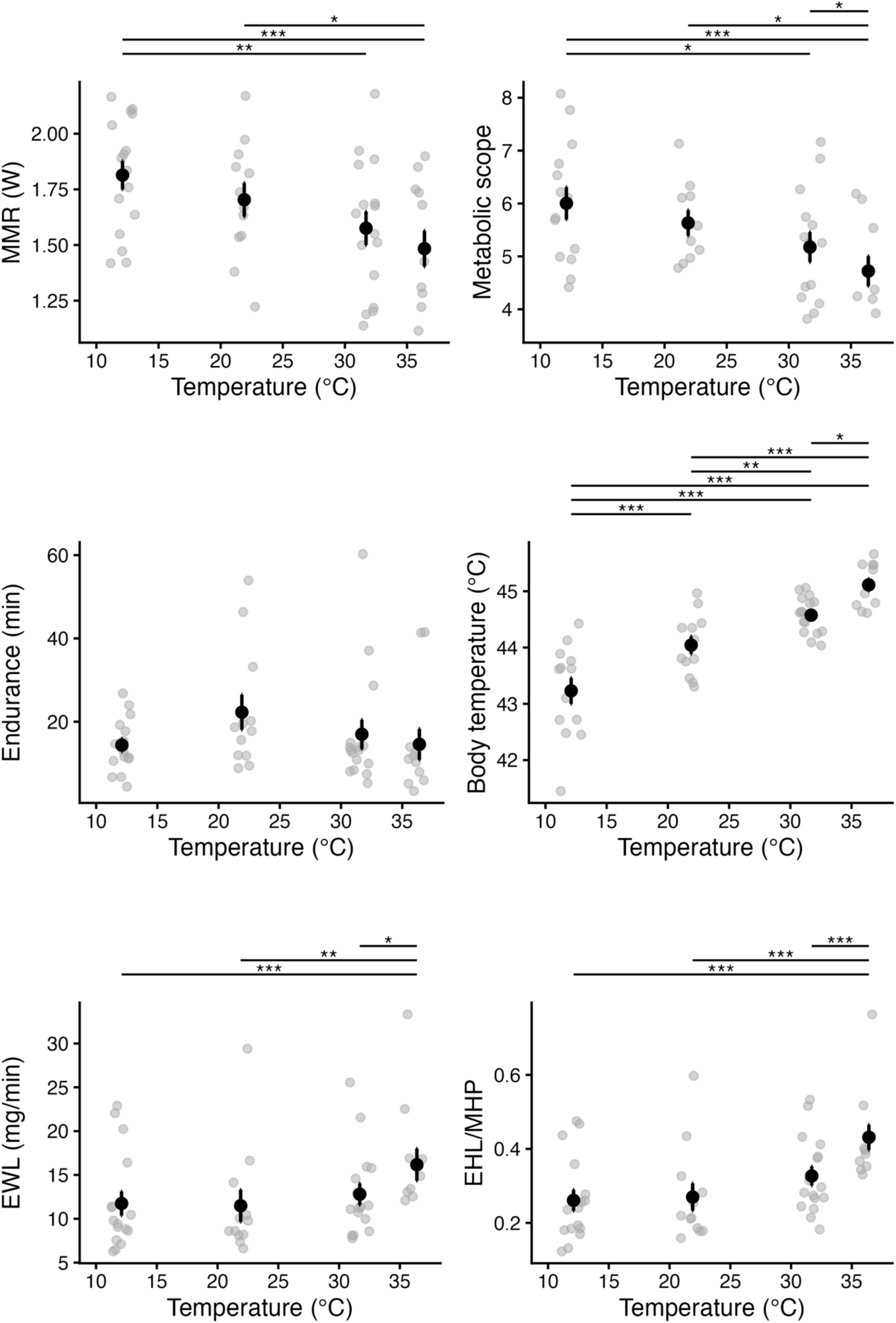
Effect of air temperature on physiological parameters during exercise. Raw data for individual is shown as gray points and mean ± se are shown as black points and error bars. Significant pairwise comparisons of estimated marginal means from a linear mixed model with cage and ID included as random factors are shown.

**Figure 3.**
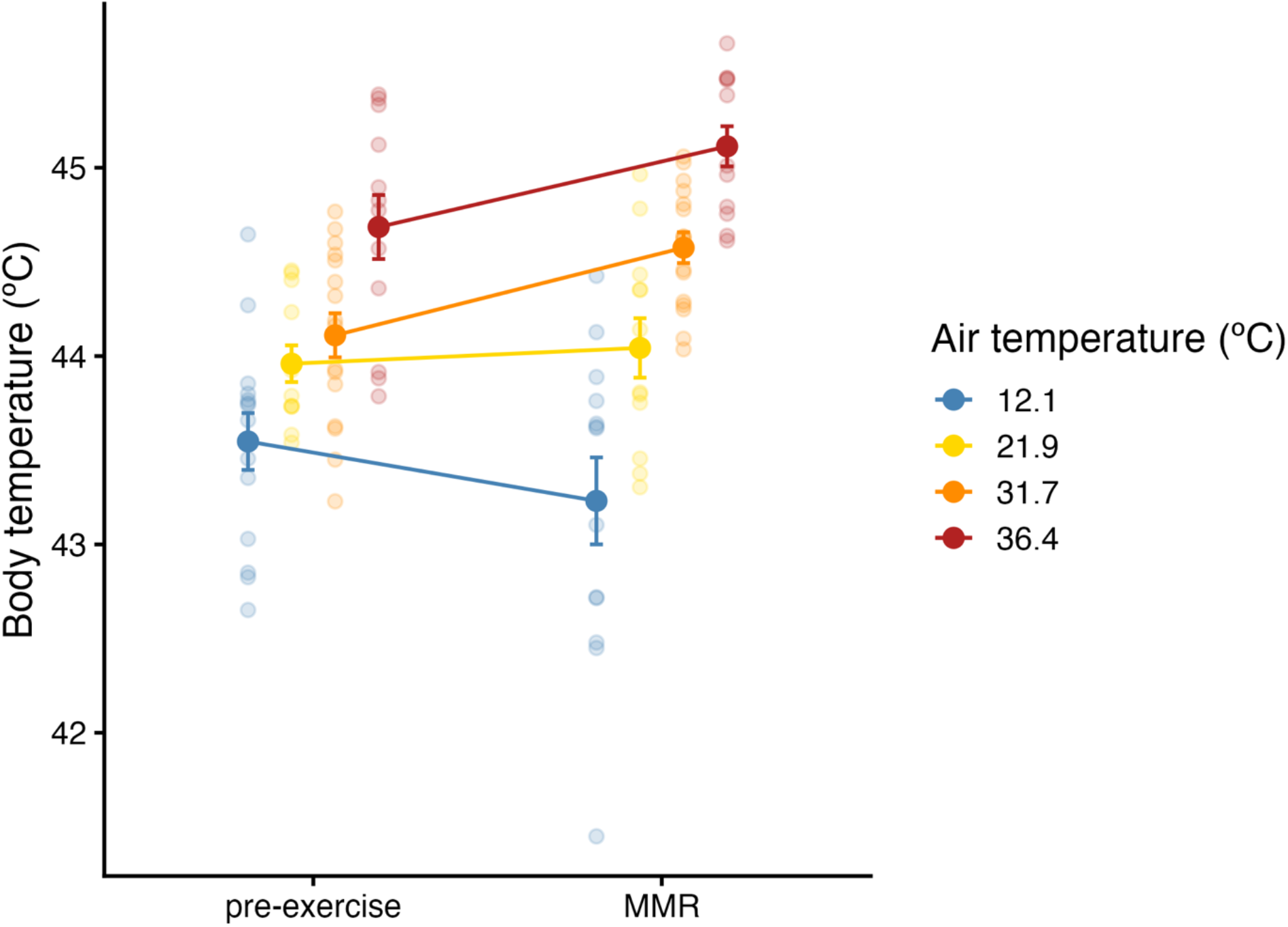
Comparison of the mean body temperature in zebra finches in the 5-minute period leading up to forced exercise (pre-exercise) and the 5-minute period when MMR was measured during exercise in four different air temperatures.

## Discussion

Our results support the hypothesis that during activity, zebra finches are prone to overheat in air temperatures well below the thermal tolerance limits measured in resting individuals. This, in turn, appeared to limit the maximum aerobic capacity and scope during activity at those temperatures (Figure 2, Table S3). In resting birds, hyperthermia occurred only at the highest temperature measured, 40 °C (Figure 1, Table S2), while in active birds, body temperature increased during exercise at both 32 °C and 36 °C (Figure 3, Table S4). During exercise in cooler temperatures, body temperature stayed stable around 44 °C (exercising in 22 °C) or decreased (in 12 °C; Figure 3, Table S4). The mean body temperature was 45 °C during activity at the highest air temperature, which has been identified as the critical upper body temperature limit in zebra finches in other studies (Cade et al., 1965; Pessato et al., 2022; Pessato et al., 2023). Therefore, birds in this study were regularly operating close to or at their upper body temperature limit.

When active in higher temperatures, the increase in evaporative cooling was seemingly insufficient to compensate for the substantial increase in metabolic heat production, reflected in the relatively low evaporative cooling capacity. The mean EWL recorded in active birds at 36 °C in this study was 0.97 g hr^-1^, which is higher than the maximum reported EWL in resting zebra finches in this and other studies of captive zebra finches (Pessato et al., 2022; Wojciechowski et al., 2021), but still lower than predicted from allometric scaling (1.19 g hr^-1^) (McKechnie et al., 2021). Mean evaporative scope was 8.22 × minimum EWL (resting within the TNZ), which is similar to predictions from allometric scaling (8.8) (McKechnie et al., 2021). However, evaporative cooling capacity (EHL/MHP), a predictor of the heat tolerance limit in dry air (Nord et al., 2026), was considerably lower (0.25 – 0.43) during exercise in this study compared to the maximum values published for resting zebra finches (1.4 in captive zebra finches, Wojciechowski et al., 2021). At most, birds lost about 5% of their body mass in water per hour, and all exercise sessions lasted less than one hour, so the birds in this study were likely not at risk of dehydration (Conradie et al., 2024). The physiological upper limit to evaporative water loss, and thus evaporative cooling capacity during exercise, likely explains why the finches became hyperthermic in higher air temperatures. When birds reach their physiological upper limits for evaporative water loss and their maximum tolerated T_b_, the only remaining avenue to balance heat gain with heat loss would be to reduce metabolic heat production, as observed in our study.

We found that there was an 18% reduction in MMR at air temperatures that are considered mild for inactive zebra finches (32 – 36 °C) compared to MMR in cooler air temperatures (12 – 22 °C). Metabolic scope decreased on average by 22% from 6.0 x BMR at 12 °C to 4.7 x BMR at 36 °C. We did not find a significant difference in endurance between air temperatures, which suggests that zebra finches can tolerate periods of increased T_b_ and EWL during activity for several minutes. However, the mean endurance was 35% higher at 22 °C (22.2 ± 14.6 min) than at both 12 °C (14.4 ± 6.3 min) and 36.4 °C (14.6 ± 12.9 min), when it was lowest suggesting that endurance could be negatively affected in both higher and lower temperatures. In humans, a main determinant of endurance during intense exercise is lactic acid threshold and accumulation (Bassett & Howley, 2000). However, exercise performance is known to decline with increasing temperatures in humans, where hyperthermia generally causes a decrease in both maximum metabolic rate and endurance due to associated changes in the function of the cardiovascular, muscular, ventilatory and central nervous system (Nybo et al., 2014). Heat stress has also been shown to decrease motor function in captive zebra finches during cognition tests (Danner et al., 2021). There is also evidence that rats cease activity when they reach a certain T_b_ threshold to prevent heat stroke (Fuller et al., 1998).

As zebra finches were forced to exercise continuously in this study, the reduction in MMR at higher temperatures may have been a direct consequence of increased body temperature (Nybo et al., 2014). However, evidence for anticipatory thermoregulation, or the preemptive reduction of performance to avoid hyperthermia, has also been found in free-living birds during nestling provisioning (Tapper et al., 2020a). We also cannot exclude the possibility that metabolic rate was upregulated in the lower temperatures due to an increased rate of heat loss during activity. These alternatives are not mutually exclusive, as birds in 12 °C could have experienced insufficient heat substitution, as evidenced by their decreasing body temperature (Table S4, Figure 3) while birds in 32 °C and 36 °C could have experienced insufficient heat dissipation.

If increased air temperature reduces the scope for activity, this could have implications for wild birds’ parental effort and reproductive success (Drent & Daan, 1980; O’Connor et al., 2022). For example, birds may be able to continue provisioning nestlings while hyperthermic, but they may not be able to maintain the same level of parental effort as they would in milder temperatures (Tapper et al., 2020a; Tapper et al., 2020b). This has been shown experimentally in wild birds with facilitated heat dissipation rates (Nord & Nilsson, 2019; Tapper et al., 2020a; Tapper et al., 2020b), and our results could potentially provide a mechanistic explanation for this pattern. This could also demonstrate a physiological link to the HDL theory in birds, which posits that energy expenditure is constrained by heat dissipation capacity (Speakman & Król, 2010). As our study was performed in artificial conditions with low forced convection compared to what would be expected during flight, with low humidity, and in the absence of solar radiation, our experimental conditions likely do not fully reflect those experienced by free-living birds. The environmental temperature where birds become hyperthermic may be shifted higher or lower in real-world conditions depending on humidity, wind or air speed, and solar radiation.

## Conclusions

As one of few studies that have measured metabolism and body temperature in exercising birds, our work improves the understanding of how heat affects aerobic capacity and aerobic scope during sustained activity. Birds in this study seemed to routinely operate near their upper T_b_ limit during activity, and were likely to become hyperthermic even in mild air temperatures. Our results show that aerobic capacity and scope were constrained by increasing air temperature due to an insufficient capacity for evaporative water loss during exercise. We believe that our findings offer a potential mechanistic explanation for how increased air temperatures could constrain parental investment, resulting in sublethal fitness costs. More research is needed to confirm a link between environmental temperature and limits to energy expenditure in free-living birds. Future studies could manipulate water availability, shade, workload, heat dissipation or a combination of these in birds while measuring activity in the wild to gain a better understanding of how heat constraints act on wild birds during activity.

## Acknowledgements

This study was supported by the Swedish Research Council (Vetenskapsrådet) to J-ÅN (2021-05467) and AN (2020-04686). ERE was supported by Stiftelsen Lars Hiertas Minne (FO2022-0336), Lunds Djurskyddsfond (50/22, 80/23, 63/24) and the Royal Physiographic Society of Lund (2023-44251).

## Data availability statement

All data and script for replication of the results, tables and figures will be available from the Dryad digital repository upon the acceptance of this manuscript.

## References

Bassett, D. R., Jr., & Howley, E. T. (2000). Limiting factors for maximum oxygen uptake and determinants of endurance performance. Med Sci Sports Exerc, 32(1), 70–84. 10.1097/00005768-200001000-00012

Bates, D., Mächler, M., Bolker, B. M., & Walker, S. C. (2015). Fitting linear mixed-effects models using lme4. Journal of Statistical Software. 10.18637/jss.v067.i01

Briga, M., & Verhulst, S. (2017). Individual variation in metabolic reaction norms over ambient temperature causes low correlation between basal and standard metabolic rate. Journal of Experimental Biology, 220(18), 3280–3289. (Journal of Experimental Biology)

Briscoe, N. J., Morris, S. D., Mathewson, P. D., Buckley, L. B., Jusup, M., Levy, O., Maclean, I. M. D., Pincebourde, S., Riddell, E. A., Roberts, J. A., Schouten, R., Sears, M. W., & Kearney, M. R. (2023). Mechanistic forecasts of species responses to climate change: The promise of biophysical ecology. Global Change Biology, 29(6), 1451–1470. 10.1111/gcb.16557

Cade, T. J., Tobin, C. A., & Gold, A. (1965). Water Economy and Metabolism of Two Estrildine Finches. Physiological Zoology, 38(1), 9–33. 10.1086/physzool.38.1.30152342

Conradie, S. R., Wolf, B. O., Cunningham, S. J., Bourne, A., van de Ven, T., Ridley, A. R., & McKechnie, A. E. (2024). Integrating fine-scale behaviour and microclimate data into biophysical models highlights the risk of lethal hyperthermia and dehydration. ECOGRAPHY, n/a(n/a), e07432. 10.1111/ecog.07432

Conradie, S. R., Woodborne, S. M., Cunningham, S. J., & McKechnie, A. E. (2019). Chronic, sublethal effects of high temperatures will cause severe declines in southern African arid-zone birds during the 21st century. Proceedings of the National Academy of Sciences, 116(28), 14065–14070. 10.1073/pnas.1821312116

Cunningham, S. J., Gardner, J. L., & Martin, R. O. (2021). Opportunity costs and the response of birds and mammals to climate warming. Frontiers in Ecology and the Environment, 19(5), 300–307. 10.1002/fee.2324

Danner, R. M., Coomes, C. M., & Derryberry, E. P. (2021). Simulated heat waves reduce cognitive and motor performance of an endotherm. Ecology and Evolution, 11(5), 2261–2272. 10.1002/ece3.7194

Drent, R. H., & Daan, S. (1980). The Prudent Parent - Energetic Adjustments in Avian Breeding. Ardea, 68(1-4), 225–252. <Go to ISI>://WOS:A1980KV89000019

Engel, S., Biebach, H., & Visser, G. H. (2006). Water and Heat Balance during Flight in the Rose-Colored Starling (Sturnus roseus). Physiological and Biochemical Zoology, 79(4), 763–774. 10.1086/504610

Engert, E. R., Nord, A., Andreasson, F., & Nilsson, J.-Å. (2026). Flexibility of exercise capacity during nestling feeding in blue tits. Journal of Experimental Biology, 229(7), jeb251043. 10.1242/jeb.251043

Fuller, A., Carter, R. N., & Mitchell, D. (1998). Brain and abdominal temperatures at fatigue in rats exercising in the heat. Journal of Applied Physiology, 84(3), 877–883. 10.1152/jappl.1998.84.3.877

King, J. R., & Farner, D. S. (1961). Energy Metabolism, Thermoregulation and Body Temperature. In A. J. Marshall (Ed.), Biology and Comparative Physiology of Birds (pp. 215–288). Academic Press. 10.1016/B978-1-4832-3143-3.50014-9

Król, E., & Speakman, J. R. (2003a). Limits to sustained energy intake VI. Energetics of lactation in laboratory mice at thermoneutrality. Journal of Experimental Biology, 206(23), 4255–4266. 10.1242/jeb.00674

Król, E., & Speakman, J. R. (2003b). Limits to sustained energy intake VII. Milk energy output in laboratory mice at thermoneutrality. Journal of Experimental Biology, 206(23), 4267–4281. 10.1242/jeb.00675

Król, E. b., Murphy, M., & Speakman, J. R. (2007). Limits to sustained energy intake. X. Effects of fur removal on reproductive performance in laboratory mice. Journal of Experimental Biology, 210(23), 4233–4243. 10.1242/jeb.009779

Lenth, R. V. (2022). emmeans: Estimated Marginal Means, aka Least-Squares Means. In (Version R package version 1.8.2) https://CRAN.R-project.org/package=emmeans

Lewden, A., Bishop, C. M., & Askew, G. N. (2023). How birds dissipate heat before, during and after flight. Journal of The Royal Society Interface, 20(209), 20230442. 10.1098/rsif.2023.0442

Lighton, J. R. B. (2018). Measuring Metabolic Rates: A Manual for Scientists. Oxford University Press. 10.1093/oso/9780198830399.001.0001

Lüdecke, D., Ben-Shachar, M. S., Patil, I., Waggoner, P., & Makowski, D. (2021). performance: An R Package for Assessment, Comparison and Testing of Statistical Models. Journal of Open Source Software, 6(60), 3139. 10.21105/joss.03139

McKechnie, A. E., Gerson, A. R., & Wolf, B. O. (2021). Thermoregulation in desert birds: scaling and phylogenetic variation in heat tolerance and evaporative cooling. Journal of Experimental Biology, 224(Suppl_1), jeb229211. 10.1242/jeb.229211

McNab, B. K. (1980). On Estimating Thermal Conductance in Endotherms. Physiological Zoology, 53(2), 145–156. 10.1086/physzool.53.2.30152577

Mitchell, D., Snelling, E. P., Hetem, R. S., Maloney, S. K., Strauss, W. M., & Fuller, A. (2018). Revisiting concepts of thermal physiology: Predicting responses of mammals to climate change. Journal of Animal Ecology, 87(4), 956–973. 10.1111/1365-2656.12818

Molenaar, E., Bebbington, K., Matson, K. D., & Kingma, S. A. (2026). Temperature-Related Changes in Avian Nestling Provisioning: A Global Analysis. Global Change Biology, 32(4), e70871. 10.1111/gcb.70871

Nakagawa, S., & Schielzeth, H. (2013). A general and simple method for obtaining R2 from generalized linear mixed-effects models. Methods in Ecology and Evolution, 4(2), 133–142. 10.1111/j.2041-210x.2012.00261.x

Nomoto, S., Rautenberg, W., & Iriki, M. (1983). Temperature regulation during exercise in the Japanese quail (Coturnix coturnix japonica). Journal of comparative physiology, 149(4), 519–525. 10.1007/BF00690011

Nord, A., Freeman, M. T., & McKechnie, A. E. (2026). Physiological constraints on heat adaptation in birds. Trends in Ecology & Evolution. 10.1016/j.tree.2026.03.006

Nord, A., & Nilsson, J. A. (2019). Heat dissipation rate constrains reproductive investment in a wild bird. Functional Ecology, 33(2), 250–259. <Go to ISI>://WOS:000458830500005

Nybo, L., Rasmussen, P., & Sawka, M. N. (2014). Performance in the Heat—Physiological Factors of Importance for Hyperthermia-Induced Fatigue. COMPREHENSIVE PHYSIOLOGY, 4(2), 657–689. 10.1002/j.2040-4603.2014.tb00554.x

O’Connor, R. S., Le Pogam, A., Young, K. G., Love, O. P., Cox, C. J., Roy, G., Robitaille, F., Elliott, K. H., Hargreaves, A. L., Choy, E. S., Gilchrist, H. G., Berteaux, D., Tam, A., & Vézina, F. (2022). Warming in the land of the midnight sun: breeding birds may suffer greater heat stress at high-versus low-Arctic sites. Proceedings of the Royal Society B: Biological Sciences, 289(1981), 20220300. 10.1098/rspb.2022.0300

Persson, E., Cuív, C. O., & Nord, A. (2024). Thermoregulatory consequences of growing up during a heatwave or a cold snap in Japanese quail. Journal of Experimental Biology, 227(2), Article jeb246876. 10.1242/jeb.246876

Pessato, A., McKechnie, A. E., & Mariette, M. M. (2022). A prenatal acoustic signal of heat affects thermoregulation capacities at adulthood in an arid-adapted bird. Scientific Reports, 12(1). 10.1038/s41598-022-09761-1

Pessato, A., Udino, E., McKechnie, A. E., Bennett, A. T. D., & Mariette, M. M. (2023). Thermal acclimatisation to heatwave conditions is rapid but sex-specific in wild zebra finches. Scientific Reports, 13(1). 10.1038/s41598-023-45291-0

Riddell, E. A., Iknayan, K. J., Wolf, B. O., Sinervo, B., & Beissinger, S. R. (2019). Cooling requirements fueled the collapse of a desert bird community from climate change. Proceedings of the National Academy of Sciences, 116(43), 21609–21615. 10.1073/pnas.1908791116

Scholander, P. F., Hock, R., Walters, V., Johnson, F., & Irving, L. (1950). Heat Regulation in Some Arctic and Tropical Mammals and Birds. Biological Bulletin, 99(2), 237–258. Doi 10.2307/1538741

Speakman, J. R., & Król, E. (2010). Maximal heat dissipation capacity and hyperthermia risk: neglected key factors in the ecology of endotherms. Journal of Animal Ecology, 79(4), 726–746. 10.1111/j.1365-2656.2010.01689.x

Tapper, S., Nocera, J. J., & Burness, G. (2020a). Experimental evidence that hyperthermia limits offspring provisioning in a temperate-breeding bird. Royal Society Open Science, 7(10). ARTN20158910.1098/rsos.201589

Tapper, S., Nocera, J. J., & Burness, G. (2020b). Heat dissipation capacity influences reproductive performance in an aerial insectivore. Journal of Experimental Biology, 223(10), jeb222232. 10.1242/jeb.222232

Torre-Bueno, J. R. (1978). Evaporative Cooling and Water Balance During Flight In Birds. Journal of Experimental Biology, 75(1), 231–236. 10.1242/jeb.75.1.231

Tucker, V. A. (1968). Respiratory Exchange and Evaporative Water Loss in the Flying Budgerigar. Journal of Experimental Biology, 48(1), 67–87. 10.1242/jeb.48.1.67

Ward, S., Rayner, J. M. V., Möller, U., Jackson, D. M., Nachtigall, W., & Speakman, J. R. (1999). Heat transfer from starlings Sturnus vulgaris during flight. Journal of Experimental Biology, 202(12), 1589–1602. 10.1242/jeb.202.12.1589

Wojciechowski, M. S., Kowalczewska, A., Colominas-Ciuró, R., & Jefimow, M. (2021). Phenotypic flexibility in heat production and heat loss in response to thermal and hydric acclimation in the zebra finch, a small arid-zone passerine. Journal of Comparative Physiology B, 191(1), 225–239. 10.1007/s00360-020-01322-0

