## supplementary material for "Air temperature limits aerobic capacity and scope in active birds"

[1] Department of Biology
Lund University
Ecology building
Lund, Sweden

[2] Swedish Centre for Impacts of Climate Extremes (climes)
Lund University
Lund, Sweden

**Supplementary Figures**

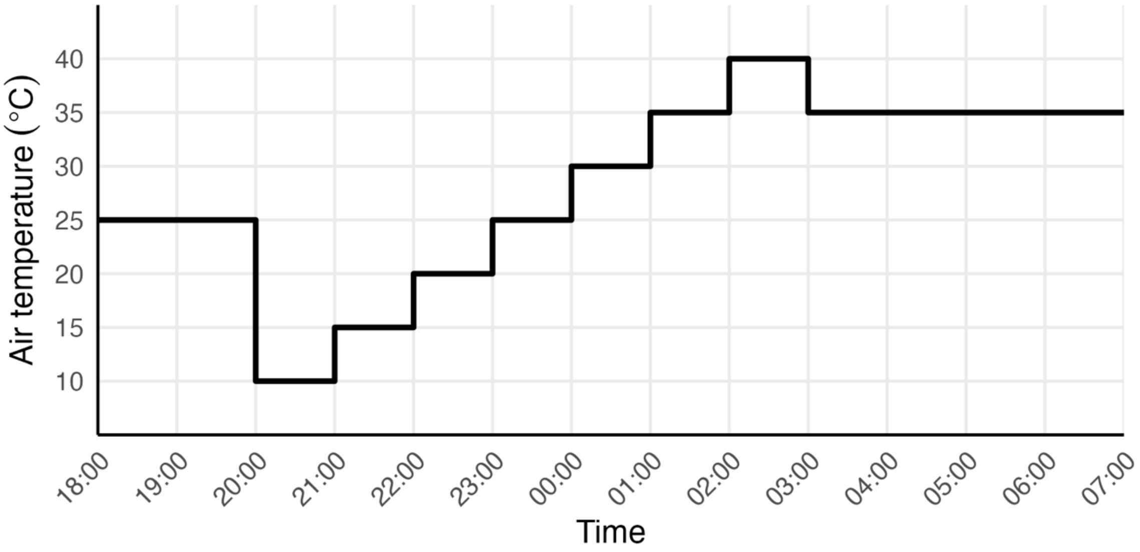


Figure S1. Heat ramp schedule for zebra finches during overnight measurements of resting metabolic rate. Birds acclimated in 25 ºC for the first 1.5 hours starting at 18:00 after which the temperature started to change to 10 ºC. BMR was measured in 35 ºC after 03:00.

### Supplementary Tables

Table S1. Sample size of zebra finches in each temperature setting for overnight measurements of RMR, and daytime measurements of MMR, and sample size for body temperature recordings (Tb) concurrent with respirometry measurements.

| Temperature | Measured temperature | SD | RMR (n) | Tb (n) |
|---|---|---|---|---|
| 10 | 10.3 | 0.9 | 26 | 21 |
| 15 | 15.5 | 1.3 | 26 | 20 |
| 20 | 20.0 | 0.8 | 26 | 21 |
| 25 | 25.1 | 0.8 | 26 | 18 |
| 30 | 29.8 | 0.7 | 26 | 23 |
| 35 | 34.8 | 0.7 | 26 | 21 |
| 40 | 39.7 | 0.8 | 26 | 23 |
| Temperature | Measured temperature | SD | MMR (n) | Tb (n) |
| 10 | 12.1 | 0.3 | 16 | 13 |
| 20 | 22.0 | 0.4 | 12 | 12 |
| 30 | 31.7 | 0.2 | 16 | 16 |
| 35 | 36.3 | 0.2 | 12 | 12 |

Table S2. Results of "Broken stick" mixed models with slopes, standard error (SE) of estimated segments and the estimated break point with 95% confidence intervals (Lower, Upper).

| Segment | Slope | SE | Break point | Lwr | Upr |
|---|---|---|---|---|---|
| RMR | | | | | |
| 1 | -0.021 | 0.001 | 33.863 | 31.727 | 38.146 |
| 2 | 0.000 | 0.005 | | | |
| Tb | | | | | |
| 1 | -0.031 | 0.007 | 33.430 | 32.761 | 33.473 |
| 2 | 0.378 | 0.017 | | | |
| EWL | | | | | |
| 1 | 0.009 | 0.003 | 36.803 | 35.908 | 37.272 |
| 2 | 0.898 | 0.076 | | | |
| EHL / MHP | | | | | |
| 1 | 0.007 | 0.000 | 36.240 | 33.768 | 36.925 |
| 2 | 0.087 | 0.006 | | | |

Table S3. Results of linear mixed-effects models (LMM) of physiological responses in zebra finches with forced exercise in four air temperatures (12 ºC, 22 ºC, 32 ºC, and 36 ºC). Response variables were maximum metabolic rate (MMR), metabolic scope (MMR/BMR), endurance, body temperature, evaporative water loss, and evaporative cooling capacity. Individual ID was included as a random effect in each model. Estimated marginal means of factors and slopes of co-variates (Estimate) or variance of random effects and residual variance, standard error (SE) of fixed effects or standard deviation (SD) of random effects and residuals, F-statistic or likelihood ratio test for random effects (Statistic) and significance (P) of model parameters and pairwise comparisons of significant factors are shown.

| Predictor | Estimate | SE (SD) | DF | Statistic | P |
|---|---|---|---|---|---|
| **MMR** | | | | | |
| Temperature | | | 34.261 | 10.450 | < 0.001 |
| 12 - 22 | 0.148 | 0.067 | 34.276 | 2.214 | 0.140 |
| 12 - 32 | 0.220 | 0.060 | 33.111 | 3.661 | 0.005 |
| 12 - 36 | 0.362 | 0.067 | 34.248 | 5.410 | < 0.001 |
| 22 - 32 | 0.073 | 0.069 | 33.695 | 1.064 | 0.714 |
| 22 - 36 | 0.214 | 0.071 | 33.595 | 3.035 | 0.023 |
| 32 - 36 | 0.141 | 0.065 | 32.894 | 2.185 | 0.149 |
| Order | | | 36.017 | 2.805 | 0.054 |
| 1 - 2 | 0.164 | 0.060 | 33.683 | 2.743 | 0.046 |
| 1 - 3 | 0.078 | 0.067 | 35.207 | 1.164 | 0.653 |
| 1 - 4 | 0.145 | 0.075 | 40.501 | 1.932 | 0.231 |
| 2 - 3 | -0.086 | 0.069 | 34.181 | -1.248 | 0.601 |
| 2 - 4 | -0.019 | 0.073 | 39.153 | -0.260 | 0.994 |
| 3 - 4 | 0.067 | 0.073 | 33.014 | 0.925 | 0.792 |
| Time | 0.012 | 0.022 | 27.463 | 0.301 | 0.588 |
| Mass | 0.052 | 0.019 | 26.262 | 7.261 | 0.012 |
| ID | 0.035 | 0.188 | 1 | 17.470 | < 0.001 |
| Residual | 0.025 | 0.158 | | | |
| **Metabolic scope** | | | | | |
| Temperature | | | 28.909 | 11.209 | < 0.001 |
| 12 - 22 | 0.615 | 0.264 | 28.860 | 2.332 | 0.114 |
| 12 - 32 | 0.721 | 0.229 | 27.241 | 3.144 | 0.020 |
| 12 - 36 | 1.485 | 0.259 | 27.357 | 5.725 | < 0.001 |
| 22 - 32 | 0.106 | 0.295 | 28.828 | 0.360 | 0.984 |

| Predictor | Estimate | SE (SD) | DF | Statistic | P |
|---|---|---|---|---|---|
| 22 - 36 | 0.869 | 0.293 | 28.489 | 2.968 | 0.029 |
| 32 - 36 | 0.763 | 0.253 | 26.451 | 3.017 | 0.027 |
| Order | | | 29.573 | 3.816 | 0.020 |
| 1 - 2 | 0.768 | 0.234 | 27.636 | 3.286 | 0.014 |
| 1 - 3 | 0.191 | 0.254 | 28.563 | 0.754 | 0.874 |
| 1 - 4 | 0.382 | 0.308 | 32.332 | 1.243 | 0.605 |
| 2 - 3 | -0.576 | 0.267 | 27.367 | -2.160 | 0.160 |
| 2 - 4 | -0.385 | 0.281 | 31.111 | -1.369 | 0.528 |
| 3 - 4 | 0.191 | 0.287 | 26.893 | 0.666 | 0.909 |
| Time | 0.115 | 0.091 | 25.404 | 1.613 | 0.216 |
| Mass | 0.118 | 0.077 | 25.748 | 2.316 | 0.140 |
| ID | 0.536 | 0.732 | 1 | 17.887 | < 0.001 |
| Residual | 0.291 | 0.540 | | | |
| **Endurance** | | | | | |
| Temperature | | | 33.438 | 2.328 | 0.092 |
| Order | | | 34.761 | 1.689 | 0.187 |
| Time | 0.145 | 0.046 | 19.320 | 10.138 | 0.005 |
| Mass | 0.058 | 0.040 | 18.720 | 2.148 | 0.159 |
| ID | 0.101 | 0.318 | 1.000 | 5.619 | 0.018 |
| Residual | 0.170 | 0.412 | | | |
| **Body Temperature** | | | | | |
| Temperature | | | 29.599 | 44.308 | < 0.001 |
| 12 - 22 | -0.759 | 0.178 | 33.894 | -4.263 | 0.001 |
| 12 - 32 | -1.460 | 0.172 | 33.477 | -8.492 | < 0.001 |
| 12 - 36 | -1.959 | 0.182 | 33.842 | -10.768 | < 0.001 |
| 22 - 32 | -0.702 | 0.173 | 31.751 | -4.047 | 0.002 |
| 22 - 36 | -1.200 | 0.178 | 31.583 | -6.745 | < 0.001 |
| 32 - 36 | -0.498 | 0.164 | 30.901 | -3.042 | 0.023 |
| Order | | | 31.373 | 5.279 | 0.005 |
| 1 - 2 | 0.626 | 0.168 | 34.135 | 3.726 | 0.004 |
| 1 - 3 | 0.422 | 0.167 | 32.663 | 2.533 | 0.073 |
| 1 - 4 | 0.379 | 0.185 | 36.629 | 2.053 | 0.188 |

| Predictor | Estimate | SE (SD) | DF | Statistic | P |
|---|---|---|---|---|---|
| 2 - 3 | -0.203 | 0.189 | 34.641 | -1.072 | 0.709 |
| 2 - 4 | -0.247 | 0.189 | 38.049 | -1.301 | 0.568 |
| 3 - 4 | -0.044 | 0.184 | 30.198 | -0.237 | 0.995 |
| Time | 0.013 | 0.048 | 19.564 | 0.073 | 0.790 |
| Mass | 0.048 | 0.041 | 17.913 | 1.346 | 0.261 |
| ID | 0.121 | 0.348 | 1.000 | 4.668 | 0.031 |
| Residual | 0.161 | 0.402 | | | |
| **EWL** | | | | | |
| Temperature | | | 36.589 | 8.544 | < 0.001 |
| 12 - 22 | -0.023 | 0.102 | 35.624 | -0.222 | 0.996 |
| 12 - 32 | -0.145 | 0.092 | 34.044 | -1.571 | 0.408 |
| 12 - 36 | -0.471 | 0.102 | 35.440 | -4.612 | < 0.001 |
| 22 - 32 | -0.123 | 0.105 | 35.522 | -1.168 | 0.651 |
| 22 - 36 | -0.448 | 0.108 | 35.048 | -4.144 | 0.001 |
| 32 - 36 | -0.326 | 0.099 | 34.255 | -3.276 | 0.012 |
| Order | | | 37.699 | 5.393 | 0.003 |
| 1 - 2 | 0.221 | 0.091 | 34.995 | 2.419 | 0.092 |
| 1 - 3 | 0.364 | 0.102 | 36.114 | 3.589 | 0.005 |
| 1 - 4 | 0.341 | 0.111 | 40.833 | 3.064 | 0.019 |
| 2 - 3 | 0.144 | 0.105 | 34.947 | 1.363 | 0.530 |
| 2 - 4 | 0.120 | 0.109 | 38.833 | 1.104 | 0.689 |
| 3 - 4 | -0.024 | 0.112 | 33.664 | -0.211 | 0.997 |
| Time | -0.086 | 0.027 | 23.520 | 10.027 | 0.004 |
| Mass | 0.023 | 0.024 | 22.894 | 0.984 | 0.332 |
| ID | 0.036 | 0.190 | 1.000 | 8.002 | 0.005 |
| Residual | 0.060 | 0.246 | | | |
| **EHL / MHP** | | | | | |
| Temperature | 0.036 | | 35.774 | 15.568 | < 0.001 |
| 12 - 22 | -0.036 | 0.032 | 36.116 | -1.132 | 0.672 |
| 12 - 32 | -0.068 | 0.029 | 34.314 | -2.354 | 0.106 |
| 12 - 36 | -0.206 | 0.032 | 35.814 | -6.508 | < 0.001 |
| 22 - 32 | -0.032 | 0.033 | 36.145 | -0.978 | 0.763 |

| Predictor | Estimate | SE (SD) | DF | Statistic | P |
|---|---|---|---|---|---|
| 22 - 36 | -0.170 | 0.034 | 35.558 | -5.073 | < 0.001 |
| 32 - 36 | -0.138 | 0.031 | 34.691 | -4.483 | < 0.001 |
| Order | -0.021 | | 36.827 | 2.928 | 0.046 |
| 1 - 2 | 0.021 | 0.028 | 35.328 | 0.729 | 0.885 |
| 1 - 3 | 0.088 | 0.031 | 36.485 | 2.799 | 0.039 |
| 1 - 4 | 0.060 | 0.034 | 40.983 | 1.764 | 0.305 |
| 2 - 3 | 0.067 | 0.033 | 35.270 | 2.061 | 0.186 |
| 2 - 4 | 0.040 | 0.034 | 38.806 | 1.183 | 0.641 |
| 3 - 4 | -0.028 | 0.035 | 33.989 | -0.797 | 0.855 |
| Time | -0.031 | 0.008 | 20.468 | 15.757 | 0.001 |
| Mass | -0.002 | 0.007 | 19.913 | 0.052 | 0.823 |
| ID | 0.003 | 0.050 | 1.000 | 4.446 | 0.035 |
| Residual | 0.006 | 0.077 | | | |

Table S4. Results of a linear mixed model (LMM) of the effect of exercise on core body temperature in different air temperatures in zebra finches. Response variables were time point (pre-exercise or MMR), temperature (12, 22, 32, and 36), and the interaction term. Individual ID was included as a random effect in the model. Estimated marginal means of factors and slopes of co-variates (Estimate) or variance of random effects and residual variance, standard error (SE) of fixed effects or standard deviation (SD) of random effects and residuals, F-statistic or likelihood ratio test for random effects (Statistic) and significance (P) of model parameters and pairwise comparisons of significant factors are shown.

| Predictor | Estimate | SE (SD) | DF | Statistic | P |
|---|---|---|---|---|---|
| Time point | | | 80.094 | 4.443 | 0.038 |
| Temperature | | | 86.768 | 53.975 | < 0.001 |
| Time point × Temperature | | | 80.095 | 5.446 | 0.002 |
| 12 | -0.311 | 0.157 | 81.601 | 1.982 | 0.051 |
| 22 | 0.084 | 0.166 | 81.328 | -0.505 | 0.615 |
| 32 | 0.465 | 0.143 | 81.328 | -3.243 | 0.002 |
| 36 | 0.428 | 0.166 | 81.328 | -2.587 | 0.011 |
| ID | 0.120 | 0.346 | 1.000 | 23.117 | < 0.001 |
| Residual | 0.164 | 0.406 | | | |
